# Protocol for the development and use of spike-in control for chromatin immunoprecipitation (ChIP) of chromatin-binding proteins

**DOI:** 10.1101/2025.05.13.653544

**Authors:** Jasbeer S. Khanduja, Mo Motamedi

## Abstract

Chromatin immunoprecipitation (ChIP) assays provide quantitative information about the genomic localization of chromatin-binding proteins. However, their sensitivity is limited by several technical variables. To generate high-confidence datasets, in this protocol, we used the *Saccharomyces cerevisiae* chromatin as an exogenous spike-in control for the ChIP of two *S. pombe* heterochromatin-associated proteins. This permitted normalization of the ChIP signals based on immunoprecipitation efficiencies across samples. Here, we describe the steps for spike-in control preparation, validation, and its use in data normalization.

For complete details on the use and execution of this protocol, please refer to Khanduja et al.^1^

## Before you begin

Chromatin immunoprecipitation (ChIP) is a powerful and widely used technique for detecting and quantifying the association of histone posttranslational modifications (PTMs) or specific proteins with different genomic regions in living cells. In this technique, cells are treated with crosslinking agents such as formaldehyde to covalently link proteins and DNA molecules in close proximity on chromatin. The immunoprecipitation (IP) of the histone PTMs or chromatin-associated protein will enrich for the DNA sequences that were in its close proximity. After ChIP, the application of next-generation sequencing (ChIP-seq) or quantitative PCR (ChIP-qPCR) on the immunoprecipitated DNA quantifies the histone PTM or protein enrichment across the genome or at a specific locus, respectively.

By performing ChIP in different genetic backgrounds or experimental conditions, ChIP assays enable a multitude of functional studies including how different chromatin-binding proteins are targeted to specific sequences in the genome, and, when combined with transcriptome analyses, how changes in the interaction of chromatin-binding proteins with the locus or genome impacts the expression of the surrounding genes. To generate robust and reproducible inferences, ChIP analyses require the generation of high-confidence datasets, permitting the detection of not only the loss or gain, but also a statistically significant decrease or increase in the ChIP signal across several samples. Considering the multi-step nature of the ChIP protocol several variables can be introduced. These variables can lead to inaccurate data interpretation, making data normalization an integral step of ChIP assays - to ensure robust, reproducible, and biologically relevant results.

Analytical and spike-in control methods are two broad approaches for ChIP data normalization which help mitigate or eliminate technical biases or variations across samples. Analytical methods rely on internal features of the data (such as input DNA control) to adjust for variability and focus on computational and statistical approaches to normalize data without introducing external experimental controls^2-5^. On the other hand, spike-in controls add a known, distinct species-specific chromatin, chromatin from engineered cells, or synthetic DNA to the samples before IP. These provide an external reference for IP normalization to reduce or mitigate IP as a variable from the ChIP protocol^6-9^. The use of spike-in controls is especially important for the detection of low-frequency DNA-protein or -PTM interactions where signal to noise ratio is generally low^7,9^. The choice of spike-in control is governed by the experimental design, target protein, and the organism in which the assay is being performed. An ideal spike-in control should have a similar chromatin structure as the experimental chromatin, its addition should not interfere with the IP of the target PTM or protein, and provide broad applicability^7,9^. Overall, spike-in controls allow for direct normalization to help reduce technical biases and support accurate cross-sample comparison under different experimental conditions.

Specifically, ChIP-Rx (ChIP with Reference exogenous genome) is a widely used and commercially available “spike-in” control for the ChIP of histone PTMs^8^ (ChIP-seq spike-in normalization reagents from Active Motif). Also, the Internal Standard Calibrated ChIP (ICeChIP) is another commercially available spike-in control for the ChIP of histone PTMs^6^, where nucleosomes reconstituted from recombinant histones and barcoded DNA serve as the spike-in control (commercially available as SNAP-ChIP from EpiCypher). These spike-in controls take advantage of the highly conserved nature of histone PTMs across eukaryotes - an antibody generated against a given PTM robustly binds to the same modification in different eukaryotic species. This means that the addition of properly prepared chromatin from other eukaryotes to each sample provides a reference exogenous genome to control for IP efficiency across samples. These spike-in controls have become widely popular in recent years and provide an easy means to generate high-confidence data for the detection of small changes in PTM signals among samples when compared with controls. Despite this, spike-in controls for the ChIP of chromatin-associated proteins are not available currently. This is mainly because chromatin-associated proteins are not highly conserved across eukaryotes, so an antibody generated against a chromatin-associated protein in one species often fails to interact robustly with its homolog in other organisms.

In recent years, the application of CRISPR technology has availed a range of new genome engineering tools for manipulating the metazoan genomes. These include the ability to add protein tags, such as FLAG, HA, GFP, GST, MYC, TAP, etc to the C- or N-terminus of the protein of interest by inserting the tag sequence into the correct sites and in-frame in the endogenous copies of the target gene. The availability of highly specific antibodies against these moieties now has provided an opportunity to develop spike-in controls for the ChIP of chromatin-associated proteins in metazoans. To demonstrate this point, recently we used the chromatin from a *S. cerevisiae* strain expressing SIR3-FLAG from a plasmid as an exogenous

FLAG spike-in control for the ChIP of FLAG-tagged heterochromatin proteins in *S. pombe*. Sir3 is a *cerevisiae*-specific protein which binds to the budding yeast silent chromatin regions. Moreover, the size of the heterochromatin in the two yeast species is roughly similar, thus providing a suitable control for our ChIP assays in *S. pombe*. By using primers against *S. cerevisiae* regions to which Sir3 binds, we were able to normalize IP efficiency across all ChIP samples. In this protocol, we provide details on the FLAG spike-in control preparation and its validation, ChIP of *S. pombe* heterochromatin proteins with the FLAG spike-in control, and ChIP-qPCR data normalization using the FLAG spike-in control.

### Cell growth, yeast cell transformation and pre-inoculum preparation

#### Timing: 8-10 days

1. Streak *S. cerevisiae* strain from a glycerol stock stored in the -80°C freezer on a yeast extract, peptone with dextrose (YPD)-agar plate and incubate the plate at 30°C for 2-3 days until well-isolated single colonies appear.
2. Select a well-isolated colony and inoculate it in 7 ml of liquid YPD media in a tube and incubate it at 30°C on a roller drum for 12-16 hours.
3. From this starter culture of *S. cerevisiae*, inoculate 100 ml of YPD media at a starting OD_600_ of 0.01, and incubate the flask at 30°C while shaking at 200 rpm.
4. At OD_600_∼ 0.6, remove the flask, spin down and wash the cells and divide them into two equal cell pellets.
5. Using a standard protocol for the transformation of *S. cerevisiae*, transform one cell pellet with the SIR-3XFLAG expression plasmid (pDM832) and the other with the empty vector (pRS315).
6. Plate the transformed cells on Yeast SD-Leu (synthetic dropout minus leucine) agar plates and incubate the plates at 30°C for 3-4 days until well-isolated single colonies appear.
7. Select a well-isolated colony for each transformed strain and inoculate in 7 ml of Yeast SD-Leu media to initiate the pre-cultures. Incubate the pre-cultures at 30°C on a roller drum for 12-16 hours.

### Formaldehyde crosslinking of the SIR3-FLAG spike-in control

#### Timing: 1 day

8. Inoculation of cultures for crosslinking
  a. Measure the OD_600_ of the pre-cultures and inoculate 400 ml of Yeast SD-Leu media at a starting OD_600_ between 0.01 and 0.02.
  b. Incubate the cultures overnight at 30°C while shaking at 200 rpm. Note: Allow for at least 6-8 cell doublings in cultures. In our hands, the *S. cerevisiae* strain transformed with a plasmid carrying the auxotrophic selection marker had a doubling time of ∼ 3 hours in SD-Leu media at 30°C. Ideally, the user should determine the doubling time of the *S. cerevisiae* strain transformed with the control or SIR3-3XFLAG plasmid and accordingly inoculate the cell cultures for crosslinking.
9. The next morning, measure the OD_600_ of the cultures. At OD_600_∼ 1.6, remove the cultures from the shaker and place them on an orbital shaker in a fume hood.
10. Crosslinking of chromatin
  a. Add 10.81 ml of 37% formaldehyde stock solution to 400 ml of the culture to achieve a final concentration of 1% formaldehyde in the cell cultures.
  b. Incubate the flasks on an orbital shaker for 15 min while shaking at a low speed. Critical: Use fresh formaldehyde from a new ampoule containing 37% formaldehyde stock. Note: Chromatin cross-linking using formaldehyde and subsequent handling of the samples containing formaldehyde should be done in the fumehood with appropriate personal protective equipment. Refer to MSDS for best handling practices and precautions.
  c. Stop the crosslinking reaction by adding 20 ml of 2.5 M glycine stock solution (final concentration of 125 mM glycine) in the cell cultures. Incubate the flasks on an orbital shaker for 5 min with shaking at a low speed.
11. Washing of cells and storage of cell pellets
  a. Transfer the cultures to two 200 ml centrifuge bottles and spin down the cells at 6000 rpm for 2 min at 4°C. Discard the supernatant in the fume hood in a biohazard container for formaldehyde disposal.
  b. Add 20 ml of ice-cold 1X TBS to each bottle and mix by vortexing. Transfer the resuspended cells to 50 ml conical tubes and spin down the cells at 4,400 rpm for 2 min at 4°C. Discard the supernatant.
  c. Wash the pellets with 20 ml of ice-cold 1X TBS as in Step 11b.
  d. Resuspend the cell pellets in 2 ml of ice-cold 1X TBS, transfer to pre-weighed and labeled 1.5 ml screw-cap conical tubes using a 1 ml micropipette.
  e. Spin down the cells at 10,000 rpm for 2 min at 4°C, remove the supernatant with a micropipette, weigh the tubes, and note the wet pellet weight.
  f. Flash-freeze the cell pellets in liquid nitrogen and store them at -80°C for future use.

### Key resources table

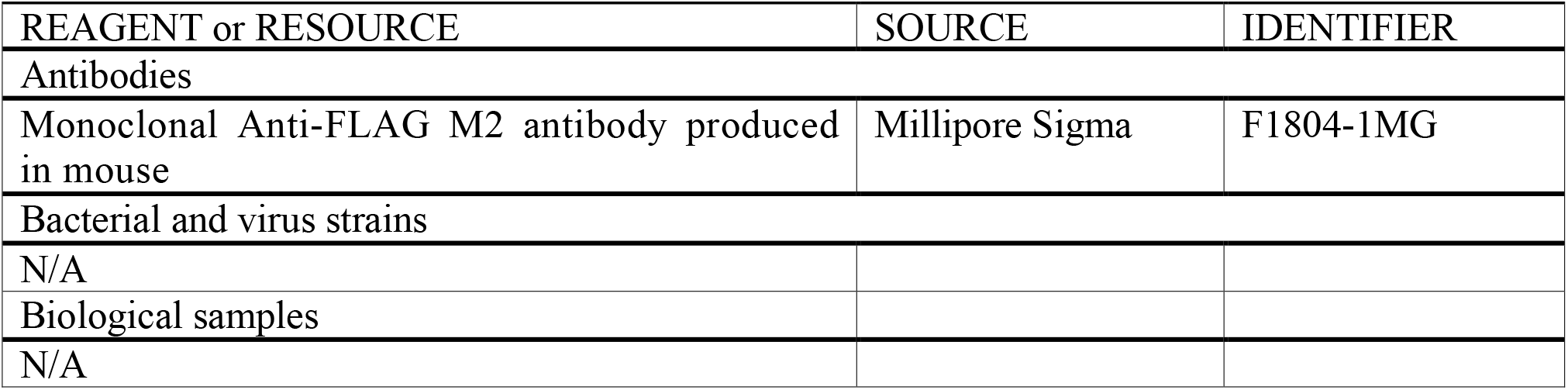
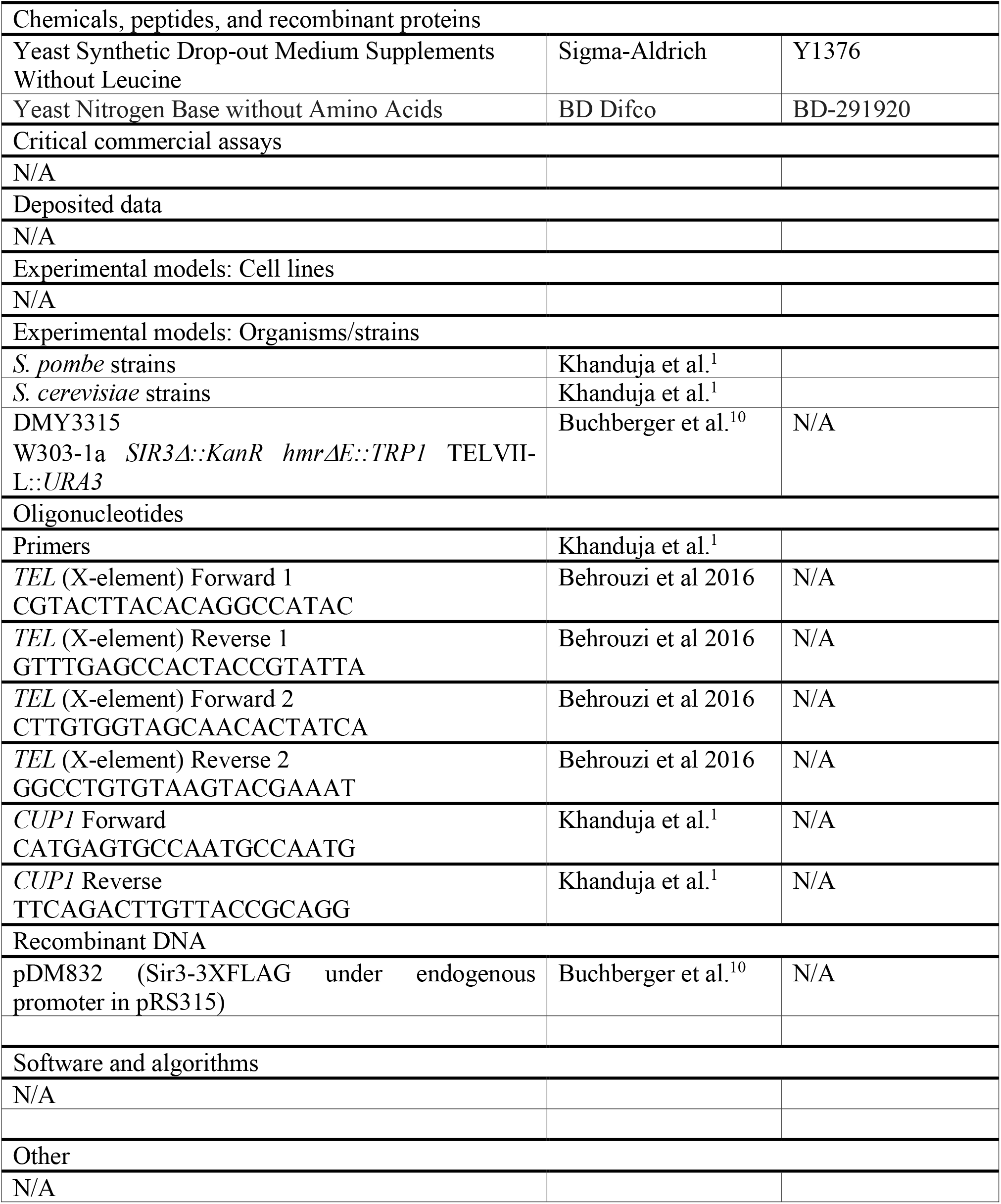

### Materials and equipment setup (optional)

For details on this please refer to the protocol published in Khanduja et al. 2025.

### Step-by-step method details

#### SIR3-3XFLAG spike-in control ChIP and its validation

<u>Troubleshooting Problems 1 and 2</u>

##### Timing: 2-3 days

In this step, SIR3-3XFLAG spike-in control is prepared. Formaldehyde crosslinked *S. cerevisiae* cell lysates were sonicated and were used to ChIP SIR3-3XFLAG using a highly specific anti-FLAG antibody.

1. Please refer to the detailed published protocol^11^ and follow steps 1-64 under “step-by-step method details” to complete the SIR3-3XFLAG ChIP.
2. Validate the ChIP using qPCR assay with primers designed to calculate % Input enrichment of SIR3-3XFLAG at its target loci (telomeres, *TEL*) and a negative control locus in euchromatin (*CUP1*).

#### ChIP of *S. Pombe* chromatin binding proteins with FLAG spike-in control

##### Timing: 2-3 days

In this step, sonicated chromatin prepared from *S. cerevisiae* cells carrying the SIR3-3XFLAG plasmid is added as the FLAG spike-in control for the ChIP of 3XFLAG-tagged heterochromatin proteins in *S. pombe*.

##### *S. pombe* cell lysate preparation, chromatin shearing by sonication, and quantification of total proteins in the cell lysate

3. Please refer to the detailed protocol published^11^ and follow steps 1-23 under “step-by-step method details” to obtain a normalized cell lysate concentration for each ChIP sample.

##### Addition of the FLAG spike-in control <u>Troubleshooting-Problem 3</u>

4. Add 150 μL of the FLAG spike-in control (chromatin prepared from *S. cerevisiae* cells carrying the SIR3-3XFLAG plasmid) to each tube. Note: We use 500 μL of normalized sonicated *S. pombe* cell lysate for each ChIP.
5. Incubate the Eppendorf tubes on a tube rotator for 5 min at 4°C in a cold room.
6. Save 2% (13 μL) of each normalized cell lysate containing the FLAG spike-in control in a pre-chilled Eppendorf DNA LoBind tube on ice and label them as Input. Store the Inputs at -20 °C until the ChIP samples are ready for crosslinking reversal. Critical: Add the same volume of FLAG spike-in control to each tube. The spike-in control should come from the same batch of chromatin prepared from *S. cerevisiae* cells carrying the SIR3-3XFLAG plasmid. Any variation in the amount, batch, or quality of spike-in control could lead to biases in data normalization.

##### Washing and pre-equilibration of Protein G Dynabeads for the ChIP assay

7. Please refer to the detailed protocol published^11^ and follow step 25 under “step-by-step method details” except that take 52 μL (13 μL for pre-clearing + 39 μL for immunoprecipitation steps) of Protein G Dynabeads/sample. Note: For 8 different samples, we will need 416 μl of Protein G Dynabeads. However, we start with Protein G Dynabeads for 9 reactions (468 μl), to account for any loss during washing and pipetting.

##### Chromatin immunoprecipitation using epitope tag-specific antibody

###### Pre-clearing

8. Pre-clear each lysate by adding 13 μL of pre-equilibrated Protein G Dynabeads and incubating the Eppendorf tubes on a tube rotator with rotation for 1 hour at 4°C in a cold room.

###### Immunoprecipitation

9. Please refer to the detailed protocol published^11^ and follow steps 26-30 under “step-by-step method details” except that in step 27 add 5.2 μg of anti-FLAG antibody to each tube and in step 30 add 39 μL of Protein G Dynabeads to each tube.

##### Washing and Elution

10. Please refer to the detailed protocol published^11^ and follow steps 31-48 under “step-by-step method details”.

Reversal of Crosslinks, Purification and ethanol precipitation of ChIP DNA

11. Please refer to the detailed protocol published^11^ and follow steps 49-64 under “step-by-step method details”.

#### ChIP-qPCR data normalization using the FLAG spike-in control

##### Timing: 1 Day

In this step, the IP efficiency of FLAG spike-in control in each sample is calculated and is used to normalize the ChIP enrichment of the *S. pombe* heterochromatin protein at the target loci in corresponding samples.

12. Use qPCR to calculate the ChIP % IP value for the spike-in control enrichment at the target locus and at a negative control locus in each sample in an experiment.
13. Calculate the normalization factor as: Normalization factor = (% IP value for the spike-in control enrichment at the target locus) – (% IP value for the spike-in control enrichment at the control locus)
14. For each sample, normalize the ChIP enrichment (in % IP) of the protein of interest at each target loci to the corresponding normalization factor calculated in step 13 above.

### Expected outcomes

A validated spike-in control in ChIP assays provides an external reference for data normalization and reproducible detection of low-level ChIP signal enrichment across different mutant backgrounds or experimental conditions. Given the modest Clr4 signal enrichment at pericentromeric repeats in cells lacking H3K9me and sRNAs, we developed a FLAG spike-in control to normalize immunoprecipitation efficiency across all ChIP samples. We first tested the efficiency of the anti-FLAG antibody to ChIP SIR3-FLAG. Consistent with a previous study^12^, we found that SIR3-FLAG was efficiently immunoprecipitated and primarily bound telomeric DNA sequences in *S. cerevisiae* (Figures 1A-1C).

**Figure 1.**
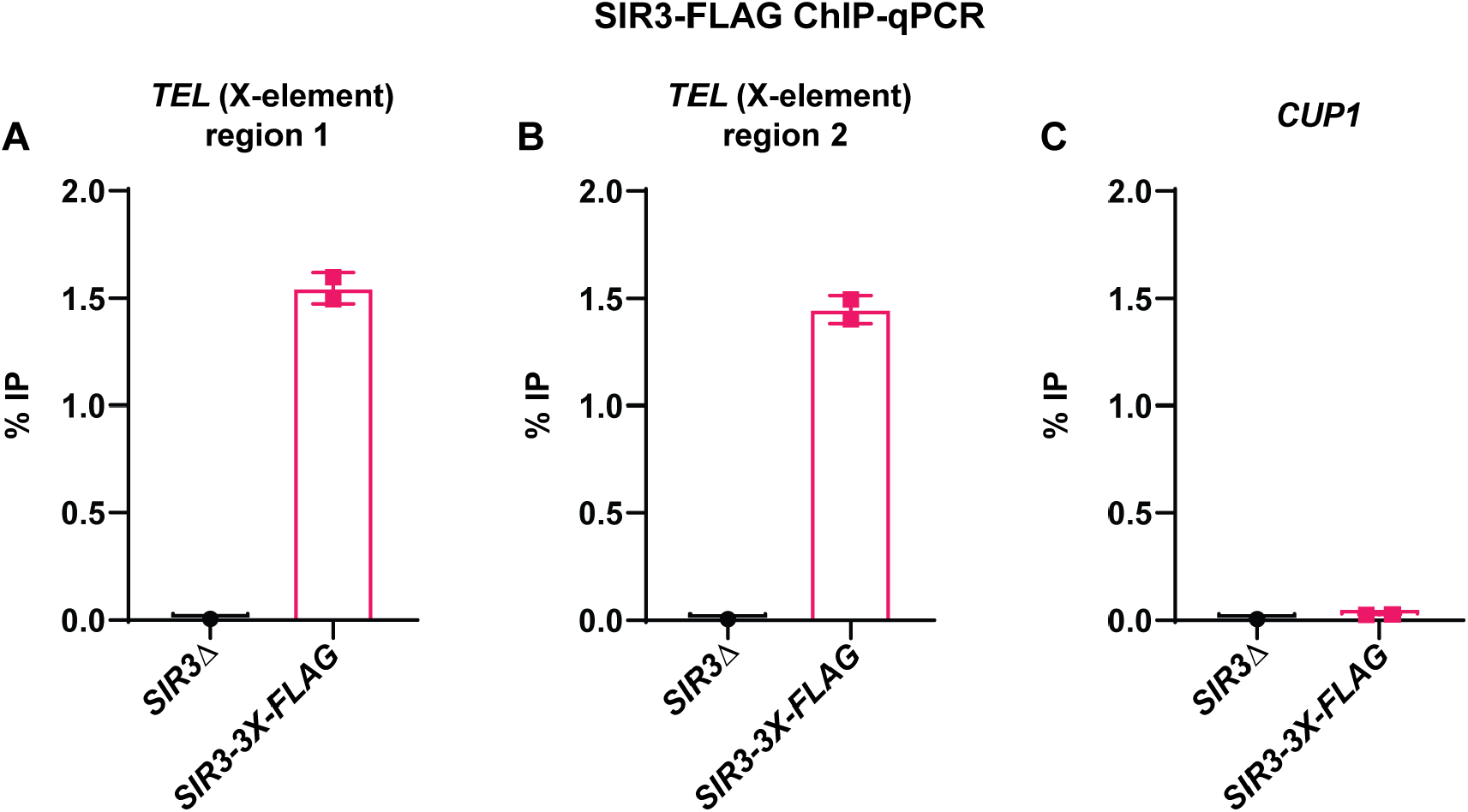
*S. cerevisiae* SIR3 is enriched at the telomeres. Graph depicting Sir3 ChIP-qPCR (mean percent IP) at *TEL* (X element) (A and B) and control *CUP1* gene (C). Error bars - S.E.; n=2 biological replicates.

Using SIR3-FLAG-containing *S. cerevisiae* chromatin as the FLAG spike-in control, we observed consistent, low-level *de novo* enrichment of Clr4 at *SPNCRNA*.*230* in cells lacking small RNAs and HDACs (*sir2Δ clr3Δ ago1Δ*) (Figure 2A). Similarly, reintroducing wild-type *clr4* or the catalytic mutant *clr4* (*clr4*.*H410D*.*C412A*) into the *clr4Δ ago1Δ* strain led to *de novo* enrichment at *SPNCRNA*.*230* (Figure 2B). In contrast, the catalytic mutant *clr4* (*clr4*.*H410D*.*C412A*) did not spread on pericentromeric repeats, unlike wild-type clr4 (Figures 2C-2E). The SIR3-FLAG-IP efficiency in the various mutant strains was comparable to that in the wild-type cells (Figure 2F). This confirmed our approach for normalizing the Clr4 ChIP signal using the FLAG spike-in control, ensuring reproducible detection of low-level Clr4 enrichment in the mutants.

**Figure 2.**
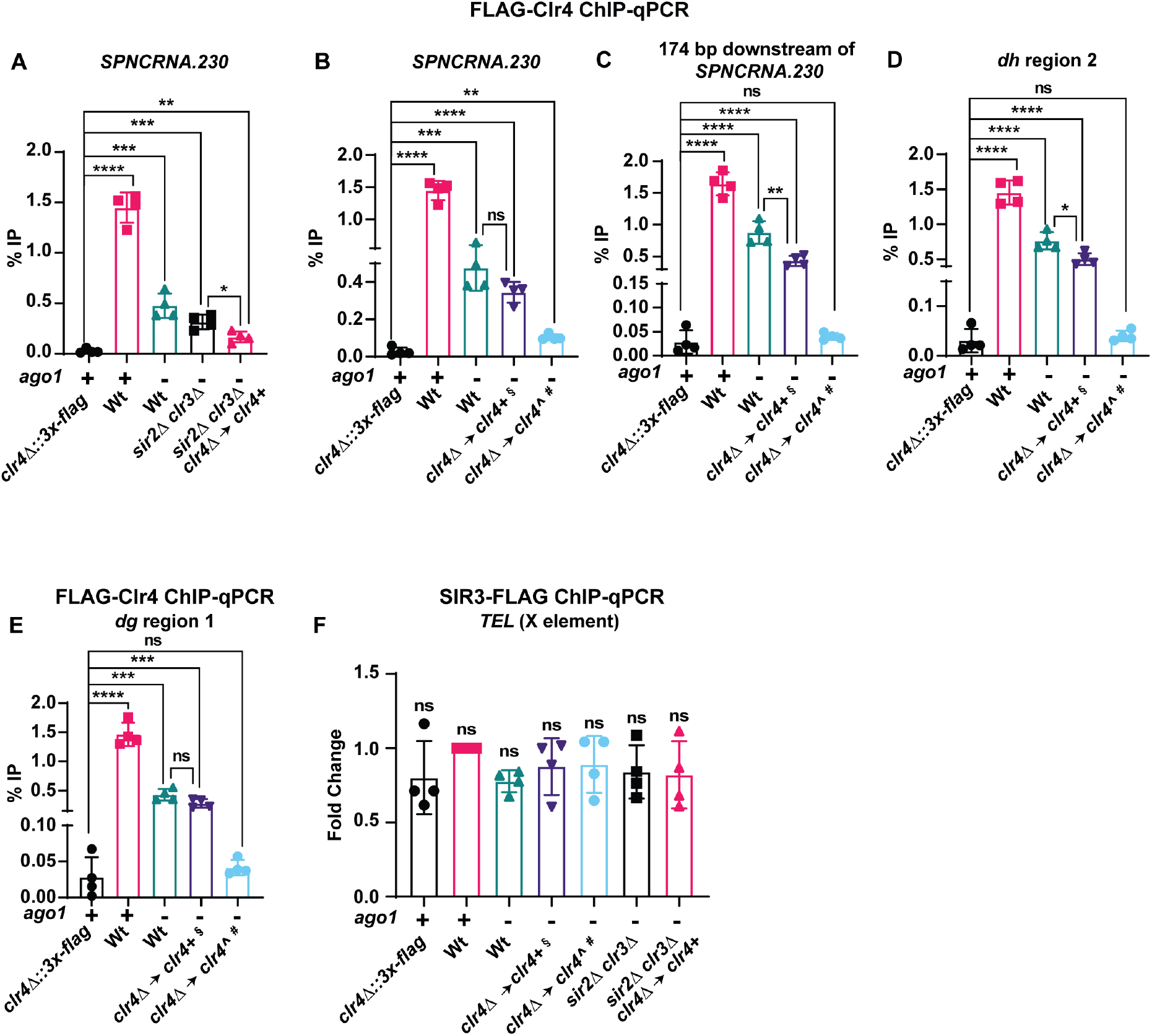
*SPNCRNA*.*230* can recruit Clr4 *de novo* and its spreading on chromatin requires its read-write activities. A) Graph depicting 3XFLAG-Clr4 ChIP-qPCR (mean percent IP) at *SPNCRNA*.*230* in the indicated strains. ‘*sir2Δ clr3Δ clr4Δ* → *clr4+*’ denotes a *sir2Δ clr3Δ ago1Δ clr4Δ* strain into which *clr4+* was reintroduced by transformation. The 3XFLAG-Clr4 ChIP-qPCR signals in the indicated strains were normalized to the *S. cerevisiae* SIR3-3XFLAG signal (FLAG spike-in control). Error bars - S.D.; n=4 biological replicates. B-E) Graphs depicting 3XFLAG-Clr4 ChIP-qPCR (mean percent IP) at *SPNCRNA*.*230* (B), 174 bp downstream from the annotated end (on the sense strand) of *SPNCRNA*.*230* (C), *dh* (D), and *dg* (E) sequences in the indicated strains. *clr4Δ::3xflag* was used as background control. The 3XFLAG-Clr4 ChIP-qPCR signals in the indicated strains were normalized to the *S. cerevisiae* SIR3-3XFLAG signal (FLAG spike-in control). Error bars - S.D.; n=4 biological replicates. F) Graphs depicting the SIR3-3XFLAG ChIP-qPCR mean fold change (percent IP) at the *TEL* X element found in the *S. cerevisiae* telomeres. The specific SIR3 enrichment at the *TEL* X element was calculated relative to the SIR3 signal at the control euchromatic *CUP1* gene. The telomeric *TEL X* element and *CUP1* gene primers do not show significant homology to the *S. pombe* genome. Error bars - S.D.; n=4 biological replicates. ^-indicates catalytically inactive version of *clr4* (*clr4*.*H410D*.*C412A*) § indicates strain in which *clr4+* was reintroduced by yeast genetic crosses. **#** indicates strain in which *clr4*.*H410D*.*C412A* allele was introduced by yeast genetic crosses. For panels A-E, Statistical significance was determined using a two-tailed unpaired Student’s *t*-test. For panel F, Statistical significance was determined using ordinary one-way ANOVA with multiple comparisons. ns *p* >0.05; \**p* <0.05; \*\**p* <0.01; \*\*\**p* <0.001; \*\*\*\**p* <0.0001 In panels A-F, data was reused for figures with permission from Khanduja J.S. et al.^1^ The data for *clr4Δ::3xflag*, wild-type, and *ago1Δ* controls in panel A have also been used in panel B.

Additionally, using the FLAG spike-in control in a ChIP of Sir2 (an H3K9 deacetylase) from hydroxyurea-synchronized cells, the normalized ChIP signal showed reproducible, low-level enrichment of Sir2 on pericentromeric repeats during the S-phase of the cell cycle (Figures 3A and 3B).

**Figure 3.**
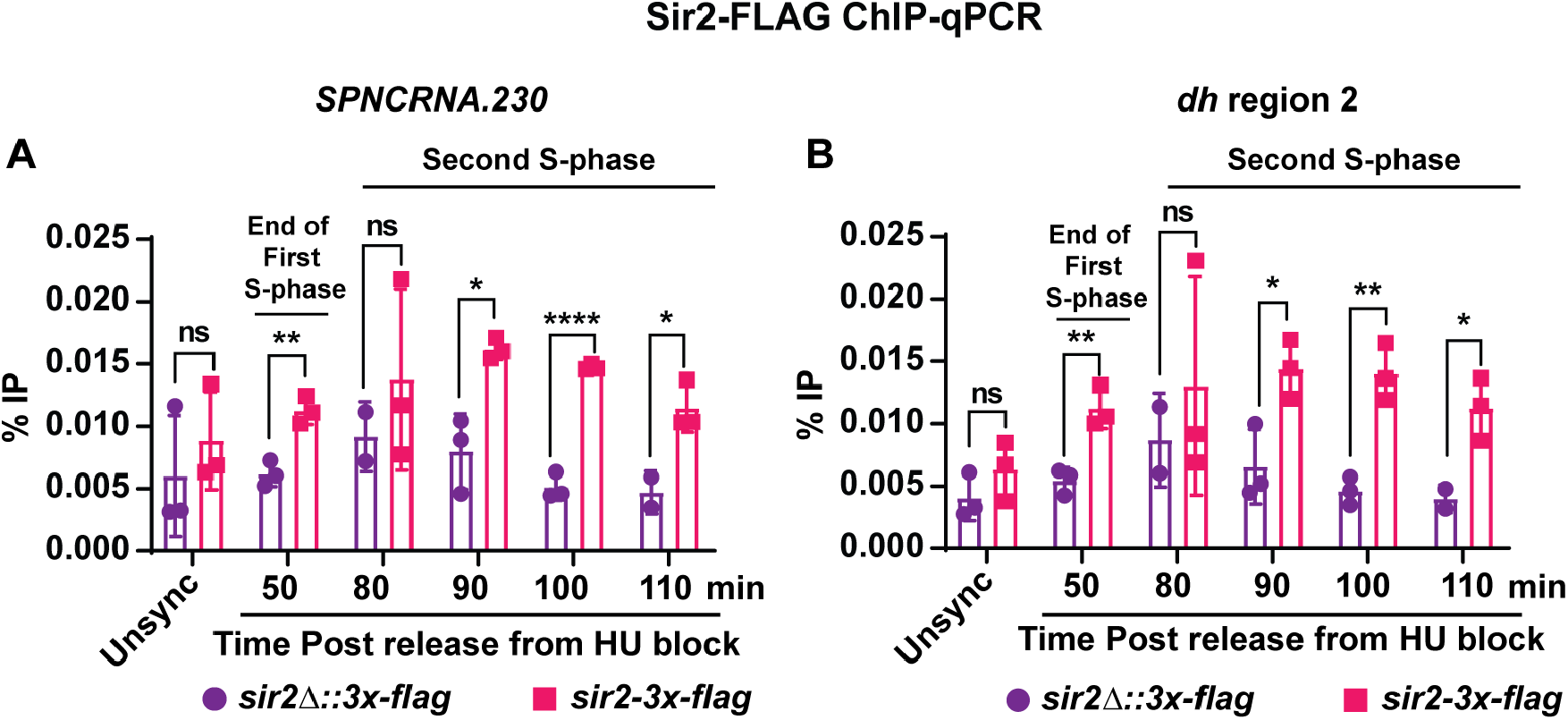
*S. pombe* Sir2 is enriched at the pericentromeres during S-phase of the cell cycle. Graph depicting Sir2-3XFLAG ChIP-qPCR (mean percent IP) at *SPNCRNA*.*230* (A) and a *dh* sequence (B) from S-phase cells synchronized by HU block and release and asynchronous cells (Unsync). *sir2Δ::3xflag* was used as FLAG background control. The Sir2-FLAG ChIP-qPCR signals in the indicated strains were normalized to the *S. cerevisiae* SIR3-3XFLAG signal (FLAG spike-in control). Error bars - S.D.; n=3 biological replicates. Statistical significance was determined using a two-tailed unpaired Student’s *t*-test. ns *p* >0.05; \**p* <0.05; \*\**p* <0.01; \*\*\*\**p* <0.0001 In panels A and B, data was reused for figures with permission from Khanduja J.S. et al.^1^

### Quantification and statistical analysis

The statistical significance of the qPCR data was determined using a two-tailed unpaired Student’s *t*-test in GraphPad Prism Version 8.3.0 (538). *n* represents the number of biological replicates and error bars show standard deviation (SD). The statistical significance was determined using GraphPad Prism Version 8.3.0, where, *p <0.05; ***p <0.001; ****p <0.0001.

### Limitations

We developed and used the FLAG spike-in control to normalize the IP efficiency in the ChIP assay of FLAG-tagged *S. Pombe* heterochromatin proteins in different mutant backgrounds, where we anticipated low to moderate enrichment of these proteins on the chromatin. This FLAG spike-in control was ideal for our study as both the SIR3 protein of the FLAG spike-in control and *S. pombe* proteins of our interest bind to heterochromatin (a repressed chromatin domain) sequences in the genome, and the size of the heterochromatin in the two evolutionary divergent yeast species is comparable.

This FLAG spike-in control may not be suitable for ChIP assay where the protein of interest in the target chromatin is not FLAG-tagged. Moreover, users should comprehensively evaluate the suitability of our FLAG spike-in control for use in the ChIP of euchromatin-associated factors.

Our protocol for cell lysis and chromatin fragmentation to make the FLAG spike-in control is identical to the protocol we use for *S. pombe* cell lysis and chromatin fragmentation for the ChIP of heterochromatin proteins of our interest. Any variation of the cell lysis and sonication procedure for preparing the FLAG spike-in control could affect its IP efficiency and, in turn, its effectiveness as a spike-in control.

### Troubleshooting

#### Problem 1

Inefficient incorporation of spike-in control in sample chromatin due to species-specific differences

The spike-in control from the chromatin of a different species may not be efficiently incorporated in the IP reaction due to differences in chromatin structure (open versus close chromatin) impacting antibody binding efficacy and consequently leading to a biased spike-in based normalization.

#### Potential Solution

Confirm that the spike-in control is efficiently co-precipitated by the antibody used in the IP step. In case of variable IP of the spike-in control, use spike-in control from a different source or use synthetic DNA spike-ins which do not interfere with IP of sample chromatin and have no chromatin structure related issues.

#### Problem 2

Inconsistent immunoprecipitation efficiency

Differences in chromatin preparation and sample handling my lead to variation in IP efficiency of test chromatin and spike-in control, which in turn, can distort the spike-in based normalization

#### Potential Solution

Optimize the IP protocol to achieve equally high IP efficiency for both the test chromatin and spike-in control in the same tube.

#### Problem 3

Inconsistent spike-in DNA quantification

Inconsistent quantification of spike-in DNA can also lead to incorrect normalization

#### Potential Solution

Ensure that the amount of spike-in control added is consistent across all the samples in an experiment. Moreover, use the spike-in control from the same batch for all the biological replicates of an experiment.

#### Problem 4

Spike-in control does not address all potential biases in ChIP data

Spike-in controls can address the biases that arise from variability in IP, and for sequencing depth in a ChIP-seq experiment. However, they do not address issues that might arise due to GC content variation, and chromatin fragmentation and accessibility biases.

#### Potential Solution

In these situations, use a comprehensive ChIP data normalization strategy that combines the use of spike-in controls and analytical methods (such as quantile normalization and GC bias correction) for robust analysis.

## Resource availability

- **Lead contact**: Further information and requests for resources and reagents should be directed to and will be fulfilled by the lead contact, Mo Motamedi.
- **Technical contact**: Technical questions on executing this protocol should be directed to and will be answered by the technical contact, Jasbeer S. Khanduja.
- **Materials availability**: This study generated new unique reagents.
- **Data and code availability**: This study did not generate new datasets or code.

## Acknowledgments

This work was supported by the National Institutes of Health grant (GM125782), American Cancer Society Research Scholar Grant (18-056-01-RMC), V Scholar grant, and Ludwig Center at Harvard grant to MM.

## Author contributions

J.S.K: protocol development, optimization, experimentation and data analysis, manuscript conceptualization, writing, editing, and final approval. M.M: supervision, funding, manuscript conceptualization, editing, and final approval.

